# Evaluating Bayesian spatial methods for modelling species distributions with clumped and restricted occurrence data

**DOI:** 10.1101/105742

**Authors:** David W. Redding, Tim C. D. Lucas, Tim Blackburn, Kate E. Jones

## Abstract

1. Statistical approaches for inferring the spatial distribution of taxa (Species Distribution Models, SDMs) commonly rely on available occurrence data, which is often non-randomly distributed and geographically restricted. Although available SDM methods address some of these problems, the errors could be more directly and accurately modelled using a spatially-explicit approach. Software to implement spatial autocorrelation terms into SDMs are now widely available, but whether such approaches for inferring SDMs are an improvement over existing methodologies is unknown.

2. Here, within a simulated environment using 1000 generated species’ ranges, we compared the performance of two commonly used non-spatial SDM methods (Maximum Entropy Modelling, MAXENT and Boosted Regression Trees, BRT) to a spatially-explicit Bayesian SDM method (Integrated Laplace Approximation, INLA), when the underlying data exhibit varying combinations of clumping and geographic restriction. Finally, we tested whether any recommended methodological settings for all methods were further impacted by spatially non-random patterns in these data.

3. Spatially-explicit INLA was the most consistently accurate method, being most or equal most accurate in 5 out of 8 data sampling scenarios. Within high-coverage sample datasets, all methods performed fairly similarly, but when sampling points were randomly spread BRT had a 1-3% greater accuracy over the other methods and when samples were clumped, spatial-INLA had a 4%-8% better in AUC score. Alternatively, when sampling points were restricted to a small section of the true range, all methods were on average 10-12% less accurate, with higher variation among the methods. None of the recommended settings for the different methods were found to be sensitive to clumping or restriction of data, except the complexity of the INLA spatial term.

4. INLA-based modelling approaches can be successfully used to account for spatial autocorrelation in an SDM context and, by taking account of random effects, produce outputs that can better elucidate the role of covariates in predicting species occurrence. Given that it is often unclear what the drivers are behind data clumping in an empirical occurrence dataset, or indeed how geographically restricted these data are, spatially-explicit INLA-based SDMs may be the better choice when modelling the spatial distribution of target species.

## INTRODUCTION

Development of quantitative methods to predict the spatial distribution of taxa from occurrence data is an active area of ecological research (Phillips, Anderson & Schapire 2006; Royle *et al*. 2012; Yackulic *et al*. 2013). Understanding where organisms are geographically located is key for many reasons: conservation scientists, for example, require knowledge about threatened species’ distributions to prioritise management efforts (Guisan *et al*. 2013). Alternatively, community ecologists need to know which species are likely to be present in the broader species pool to better understand the community assemblage process at a specific location (Pellissier *et al*. 2010; Thuiller *et al*. 2015). Within disease research, knowledge about pathogen distributions across a landscape can better inform understanding of spatial patterns of human disease risk (Peterson 2006; Redding *et al*. 2016).

Statistical approaches for inferring the spatial distributions of taxa across landscapes are commonly termed ‘Species Distribution Modelling’ (SDM) or ‘Niche Modelling’. Rather than estimating niches *per se* or looking to create models to better understand the causative process behind spatial distributions, in most cases these statistical approaches are used as a spatial interpolation across a region of interest to overcome incomplete sampling and predict the probability of presence/absence at all unsurveyed locations. SDMs commonly rely on regression techniques which identify the correlative associations of species’ occurrence to a suite of explanatory and spatially extensive variables, e.g. temperature, altitude, and rainfall (Phillips, Anderson & Schapire 2006; Elith *et al*. 2011).

Over the last decade there has been a significant uptake in methods that fit highly complex SDM models, for example using maximum-entropy based lasso regressions or boosted-regression tree (BRT) approaches (Elith *et al*. 2006). This has been driven by the ability of these methods to quickly produce models with high ‘in-bag’ predictive accuracy, as measured using, for instance, the ‘area under receiver operating curve’ (AUC) statistic (Qiao, Soberón & Peterson 2015). The availability of bespoke software packages such as MAXENT (Phillips, Anderson & Schapire 2006) and the R package ‘dismo’ (Hijmans 2013) have helped augment this methodological uptake, as these packages are computationally inexpensive, user-friendly, free-to-user and produce visually appealing outputs.

The ease of use of these packages, however, can also drive a method-agnostic (or ‘standard settings’) approach to analysis (Yackulic *et al*. 2013). By default, highly complex, ‘black box’ approaches can over-fit models to the data so that any predictions partially reflect the sampling biases of input datasets (Elith *et al*. 2006; Phillips, Anderson & Schapire 2006). Various methodological configurations have been suggested to prevent such problems arising, including: reducing the complexity of the final model by increasing penalties on additional parameters (Anderson & Gonzalez 2011), reducing the number of predictors (Syfert, Smith & Coomes 2013), accounting for the spatial patterns of samples by using background points generated with a similar spatial structure (Warton, Renner & Ramp 2013; Beck *et al*. 2014; Stolar & Nielsen 2015), or reducing the spatial autocorrelation of the sampling points in the analyses (Miller 2012; Record *et al*. 2013; Crase *et al*. 2014).

A straightforward approach to control one aspect of sampling bias, i.e. the spatial patterning of samples, is to directly incorporate a spatially-structured random term into the model. However, until recently formally incorporating such a term into many commonly used SDM frameworks was prohibitively complex (Record *et al*. 2013). Integrated Laplace Approximation (INLA) Bayesian additive models have been recently implemented in the programming language R (Lindgren & Rue 2015), offering a highly flexible modelling environment which can incorporate a variety of spatial effects into binomial presence/absence models. INLA methods analytically determine the posterior marginal distributions for parameters (Beguin *et al*. 2012), which affords a large reduction in computational time compared to ‘search’ based (e.g. MCMC) methods. This approach results in fast Bayesian models that have been shown to be effective in producing SDM-type spatial predictions (Beguin *et al*. 2012), and can additively incorporate many different complex terms to account for ‘random’ effects, such as spatial, observer and sampling bias.

It is unclear, however, whether Bayesian methods that remove the effects of spatial autocorrelation are needed when inferring SDMs, and in what situations they might offer an improvement over existing approaches. Empirically collected samples are known to be often spatially biased due to anthropogenic drivers, such as: Spatially heterogeneous reporting rates around areas with high numbers of observers and biases towards areas with active sampling programmes (Hortal *et al*. 2008; Boakes *et al*. 2010), or, high disease detection rates where there are identification/diagnostic facilities (Bern, Maguire & Alvar 2008). However, some or all of the spatial patterning contained within any set of taxonomic samples could also be caused by the underlying suitability of the environment, rather than the impact of anthropogenic drivers, meaning the clumping of points itself would contain important information and should not be discounted. Here, therefore, we test the role that clumping and geographical bias have on the predictive ability of presence-background modelling methods, comparing a spatially-explicit Bayesian approach (INLA) to two common, non-spatial methods (MAXENT and BRT) on sets of simulated data that show high variation in clumping and bias. We also test, for all methods, if any of the ‘best-practice’ user settings previously recommended for optimum SDM analysis are sensitive to our measured data biases. Overall, we show how the processes behind the spatial patterns of the input data can dictate the optimal choice of methods, showing that for the commonly encountered scenario of having clumped sample data, spatial Bayesian models are more consistently accurate than traditional methods.

## METHODS

We tested the predictive performance of different SDM methods to reconstruct spatial presence, using simulated data sets with varying degrees of data clumping and geographical bias for 1000 hypothetical taxa. All analysis was performed using R (R Development Core Team 2015). For each taxon, we generated four sets of simulated data with which to reconstruct occurrence, as follows:

1. *Covariate raster layers*. We first generated 5 random covariates (to represent bio-climatic data layers) across a hypothetical landscape of 1 degree grid cells covering the world (a common resolution for SDM models and input data) using two-dimensional vertical, horizontal or diagonally ‘sloped’, or, ‘bell-shaped’ gradients for each covariate across the full extent of the landscape.
2. *True presence raster layer*. We employed the covariates in a presence-absence binomial regression, with randomly generated slopes and intercept, to predict the true spatial distribution of each hypothetical taxon. The regression formula was generated using a random number of terms taken from a pool consisting of all linear and square terms for each covariate and first order interactions between each of the five terms, giving a total of 25 possible terms in the most complex models. We generated the ‘true presence’ layer by then spatially predicting the generated regression model using the original covariate raster layers as inputs. After generating this layer, we added a small amount of random noise to each grid cell (jitter function; R Development Core Team 2015) to create a more realistic problem, where exact covariates relationships are unknown (due to uncertainty in remotely-sensed data, for example).
3. *Validation data*. We sampled the true presence layer using 1000 random locations to create a validation data set of true presence and absence points, which we could use to evaluate the predictions from different SDM methods.
4. *Spatially biased sample data*. We then created a ‘samples’ data set with differing amounts of clumping and geographical bias, analogous to a typical input data used in an empirical SDM analysis. To create the biases, we first selected a small random number of ‘seed’ points from across the simulation landscape. These seed points were then filtered (i.e. removed or not) using a spatially heterogeneous ‘sample reporting’ layer reflecting that, at some locations, external factors would limit reporting of possible records. This reporting rate varied randomly between 0 and 1 within six, randomly-sized and placed, but contiguous, areas across the landscape. We then generated the final SDM input dataset by randomly drawing a set of ‘sampling’ points from around each remaining ‘seed’ points, with a random clumping coefficient (the mean of a Gaussian distribution, ranging from 1-50 with a standard deviation equal to the mean divided by 5) dictating how tightly clustered any secondary sampling points were around each seed point. The number of ‘samples’ around each ‘seed’ was constrained to be either conditionally dependent on the probability of true presence or unaffected by suitability of the landscape. We term these two processes ‘biological’ bias and ‘random’, respectively. Therefore, for random datasets, sampling density was entirely random with respect to the underlying habitat suitability, representing the situation where any spatially heterogeneous sampling effort is driven by convenience or other non-biological processes. For biological bias, the density patterns in the spatially-biased samples were driven by the probability of the taxon’s true presence, representing the situation where a greater number of reports are made where there are more actual individuals to observe. All the secondary points (ranging between 25 and 500 points) were then used as the final sampling dataset to be brought forward to the SDM analysis.
5. *Analysis*. After simulation, we measured two aspects of the spatial pattern of the sample points in such a way that could be applied to a typical SDM input dataset, as follows: Clumping was defined as Clark-Evan’s dispersion coefficient of the samples (Baddeley, Rubak & Turner 2015) and split into high and low categories (clumped & even) by the median value; Geographical bias was calculated as the area covered by a convex hull containing all the biased samples divided by the range of occurrence of the true positive (validation) samples, again split into high and low categories (high coverage & restricted) using the median value. This latter measure represents when the state of knowledge, for a given species, is solely the distributional limits of its geographical range of occurrence. Each dataset was then assigned to one of four equal groups, termed: even & high coverage, clumped & high coverage, even & restricted, and clumped & restricted. For R code, see supplementary code S1.

For each taxon, we reconstructed the geographic range of occurrence using the simulated sampling data set and the covariate raster layers using the following SDM methods: (1) Maximum Entropy (MAXENT, Phillips *et al*. 2004); (2) Boosted Regression Trees (BRT, Elith 2009); (3) a Bayesian additive spatially-explicit modelling approach with Integrated Laplace Approximation, INLA (Rue, Martino & Chopin 2009); and (4) a non-spatially explicit INLA Bayesian model (to ascertain whether the performance of previous INLA model (3) was due to the spatial random effect or other aspects of the INLA model). We then generated predictions for the probability of presence, for each species, at each of the locations of the 1000 validation data points, and used an AUC (Area Under operating Curve score) approach to calculate the predictive accuracy of each method by comparing the validation data with the predicted presence value. AUC represents a calculation of predictive accuracy across all possible classification threshold values, such that a value below 0.5 indicates a predictive ability equal to random expectation and 1 perfect predictive ability (Qiao, Soberón & Peterson 2015).

Finally, we tested whether any ‘best-practice’ configurations for setting up any of the methods were sensitive to clumping or geographical restriction of the data. We employed a brute-force approach, testing all reasonable SDM method configurations (see Table 1) and ensuring that we included those previously identified method set-ups shown to improve predictive accuracy.

**Table 1.** Options tested for four different species distribution model methods (for details see text). Numbers in 'Setting' columns indicate known precedence in the literature: 1 = (Merow *et al*. 2014), 2 =(Phillips *et al*. 2009), 3 = (Syfert, Smith & Coomes 2013), 4 =(Beguin *et al*. 2012), 5=(Record *et al*. 2013), 6=(Elith, Kearney & Phillips 2010), 7 = (Fourcade *et al*. 2014).

| METHOD | SETTING | OPTIONS |
| --- | --- | --- |
| MAXENT | Beta – regularization (BR)<sup>1</sup> | Value of 1 to 20 |
| MAXENT | Model complexity (MC)<sup>1</sup> | Any combination of: nolinear, noquadratic, noproduct, nothreshold, nohinge, noautofeature |
| SPATIAL<br>INLA | Mesh complexity (CO)<sup>4</sup> | Cut off from 0.5 to 15 |
| BRT | Bag Fraction (BF)<sup>6</sup> | 0.5, 0.65, 0.75 |
| BRT | Learning rate (LR)<sup>6</sup> | 0.0001 to 0.1 |
| BRT | Tree complexity (TC)<sup>6</sup> | 2 to 8 |
| BRT | No. of trees for prediction (NT)<sup>6</sup> | First 200, first 400, Best guess |
| ALL | Covariate choice (CC)<sup>3</sup> | Most parsimonious model chosen by minimal AIC |
| ALL | Spatial thinning (ST)<sup>7</sup> | Spatially-thinned presence points to reduce autocorrelation |
| ALL | Spatial weighting of pseudo-absence (SW)<sup>2</sup> | Spatially-correlated pseudo-absence pts |
| ALL | Random pseudo-absence (R)<sup>2</sup> | Random pseudo-absence pts |

## RESULTS

When comparing across sampling bias scenarios, spatial INLA was the most consistently accurate, being either the most accurate, equal most accurate or second most accurate method in 5 out of the 8 combinations examined here (Figure 1). The proportion of the simulated landscape covered by the sampling points was a key factor in dictating predictive accuracy, though presence of clumping did also confer a small loss in accuracy (Figure 1). Within the high coverage scenarios, when analysing datasets with low sample clumping (even & high coverage) all methods gave high predictive performance (Figure 1), but with BRT the most accurate (mean AUC 0.955 and 0.935 for biological and random clumping processes respectively), and with non-spatial INLA the least accurate (0.93 and 0.89 for biological and random clumping). For high coverage datasets with significant clumping of points (clumped & high coverage), the most accurate method was the spatially-explicit INLA model for both biological and random underlying processes (mean 0.929 and 0.901 AUC), again with the non-spatial INLA performing the poorest (mean 0.914 and 0.865 AUC) (Figure 1).

**Figure 1.**
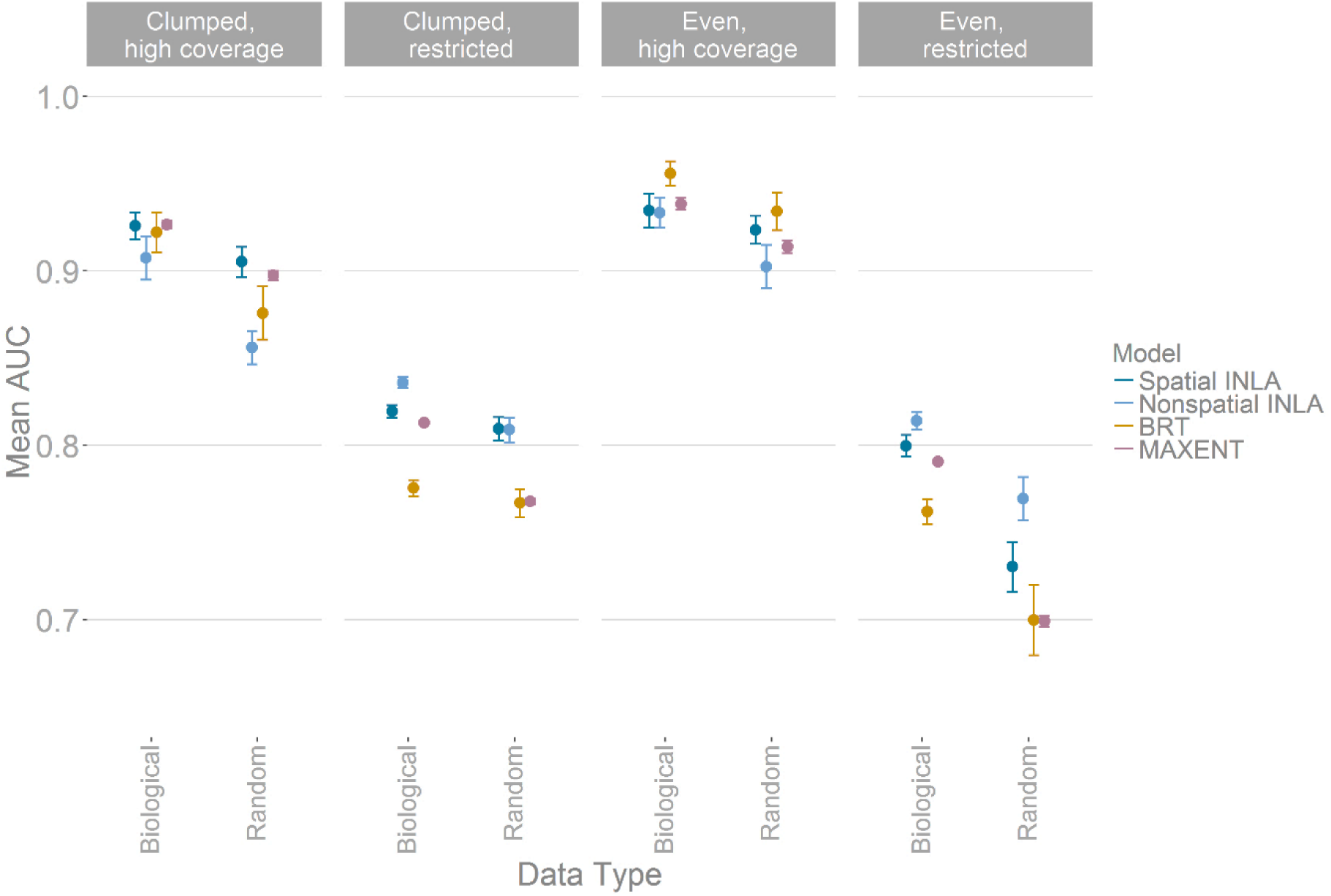
Comparison of the mean accuracy (AUC) of SDM models over 1000 simulatedtaxa. Sample clumping is caused by either biological bias or random processes. Panels show the predictive accuracy of data subsets binned into either high or low clumping and high or low coverage of the simulated “true” range. Points represent mean AUC scores from 1000 validation points per taxa and whiskers 95% confidence intervals around each mean, where scores less than 0.5 represent no accuracy gain over random chance. Spatial INLA – Bayesian SDM inferred using Integrated Nested Laplace Approximation with a spatial autocorrelation term, Non-spatial INLA – Bayesian SDM inferred using INLA without a spatial autocorrelation term, BRT – boosted regression trees based SDM, MAXENT – maximum entropy based SDM.

Within low coverage sampling datasets (even & restricted and clumped & restricted), there were similar among method patterns, with an average low predictive accuracy compared to high coverage sampling data (0.71-0.84 AUC scores across all methods) and higher variance in scores. In both cases, the simplest modelling approach, non-spatial INLA models, tended to perform better (Figure 1), with spatial-INLA the next most accurate method. When comparing clumping processes, if clumping was driven by biological processes rather than by random processes predictive accuracy was generally higher (Figure 1).

For most methods, there was no difference in terms of predictive accuracy when choosing the best set-up of analysis options across clumped and restricted sampling groups. For instance, in all cases, randomly placed pseudo-absences/background points (R - Table 1 & Figure 2) out-performed both spatially-weighted absence points (SW) and spatially-thinned presence points (ST) in our analysis (Figure 2). Reducing the number of covariates always reduced average predictive accuracy (average reduction of 0.068 AUC score across methods) and including interaction terms in the formulas resulted in no significant gains in accuracy, irrespective of the complexity of the function used to generate the simulated data (Figure S1).

**Figure 2.**
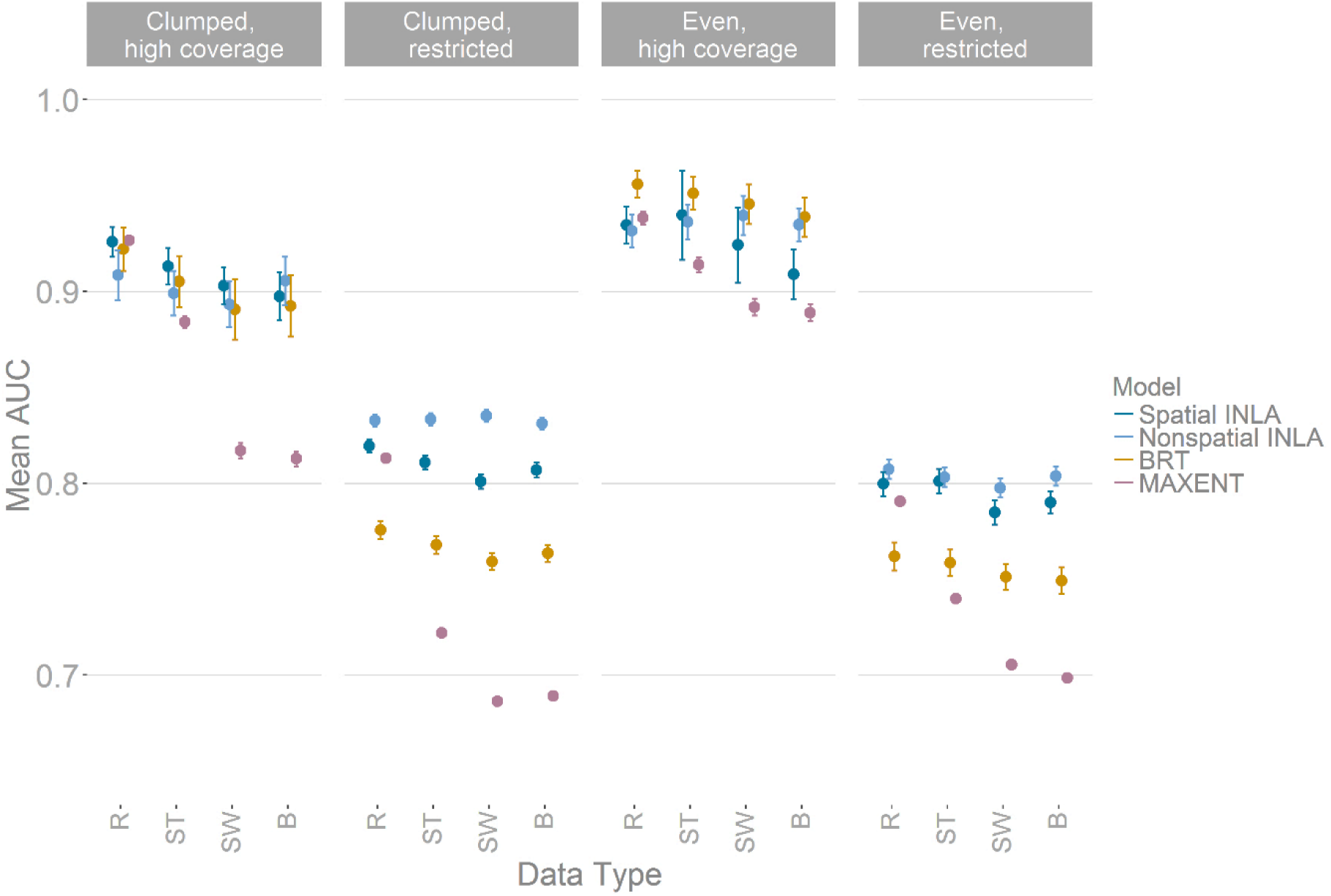
Comparison of the mean accuracy (AUC) of SDM models over 1000 simulated taxa when altering the pseudo-absence (background) point configurations and the effects of spatial thinning of presence points, on four SDM methods and across 4 types of dataset with different clumping and spatial bias. Panels show the predictive accuracy of data subsets binned into either high or low clumping and high or low coverage of the simulated “true” range. Where R represents random absence points, ST – spatial thinning, SW – spatially weighted absence points, B – both weighting and thinning (Table 1) and Spatial INLA – Bayesian INLA model with spatial autocorrelation term, Non-spatial INLA – Bayesian INLA model without spatial autocorrelation term, BRT – boosted regression trees, and MAXENT – Maximum entropy based model. Points represent mean AUC scores from 1000 validation points per taxa and whiskers 95% confidence intervals around each mean, where scores less than 0.5 represent no accuracy gain over random chance.

For INLA models comparing cut-off values from 0.5 to 8, we show that for high coverage-high clumping datasets (Figure 3) the smaller the cut off (and therefore the more complex the resulting spatial term), the more accurate the final models are. For data with low clustering and high coverage and with both low coverage datasets (Figure 3), there appears to be the opposite relationship with highest accuracy at values at a cut-off of around 3 to 6. For MAXENT models, increasing the regularisation (beta) setting, which preferentially selects less-complex models, generally resulted in less accurate results (Figure S2). Also, when manually specifying model ‘features’, the inclusion of ‘hinge’ factors was important for good predictive accuracy (Figure S3). None of the BRT initial set-up values (Figure S4-S7) produced any clear difference in predictive accuracy.

**Figure 3.**
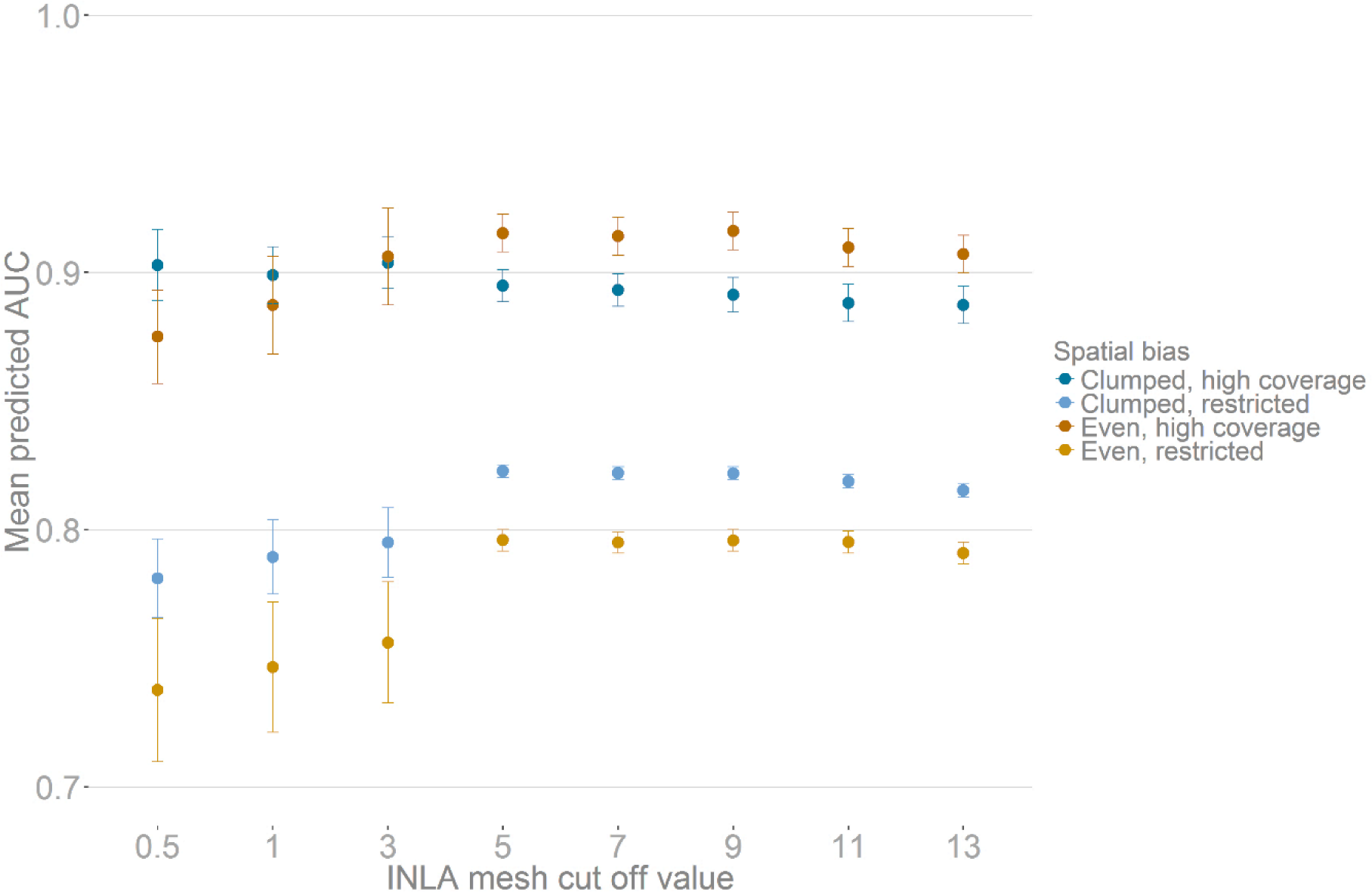
Comparison of mean accuracy (AUC) of spatial INLA-based SDM models on 1000simulated taxa when varying the complexity of the underlying spatial mesh. Colours show the predictive accuracy of data subsets binned into either high or low clumping and high or low coverage of the simulated “true” range. Points represent mean AUC scores across 1000 taxa and whiskers 95% confidence intervals around each mean, where scores less than 0.5 represent no accuracy gain over random chance.

## DISCUSSION

Choosing the best method to undertake species distribution modelling depends on the spatial patterning within the data. For instance, on even & high coverage datasets all methods performed well, especially boosted-regression trees. Such ‘ideal’ datasets are likely to be infrequently found but when they are, the ease of use and high accuracy of MAXENT and BRT mean these methods make them ideal choices. For all types of clumped data, the spatial INLA Bayesian model performed consistently well by assuming that any spatial patterning is noise and relying instead on covariate relationships.

It is expected, and supported here, that clumping driven by underlying suitability has less of an impact on the predictive performance of the models than random clumping, but, importantly, not by a large amount. In the case where SDMs are inferred using data sources such as GBIF, the complex processes underlying any spatial patterns are often unclear (Beck *et al*. 2014), and it appears that a precautionary approach of using a spatially-explicit method would reduce the impacts of misidentifying the processes behind underlying spatial bias. Sampling restriction, however, had a much larger impact on predictive accuracy than clumping. Building a SDM for a whole species based on sampling just one part of the range risks biasing the underlying regression models, as opposed to clumping which will more likely just add ‘noise’. Here, the simple models produced by non-spatial INLA (and likely GLM approaches) appear to do well, perhaps because they are less ‘fitted’ to any biases in the data, producing more general predictions.

Our results, therefore, show that it is important to remain sceptical about SDM predictions with high ‘in-bag’ AUC scores if there is no explicit measure of the proportion of the known range that has been sampled. With restricted samples, within our study, results with ‘in-bag’ AUC scores >0.95 in our simulations often had a real AUC score of less than 0.75. Overall, without taking into account the data sampling scenarios examined here, and instead evaluating SDM methods using ‘ideal’ (even & high coverage) datasets, most analyses would prefer a BRT approach, which was a poor performer on the clumped and restricted datasets.

In terms of evaluating ‘best-practice’ settings for each method, irrespective of the data sampling scenario, reducing the number of covariates to decrease collinearity was again shown to be an unuseful approach (Syfert, Smith & Coomes 2013). It seems that even variables with a minimal ability to explain the variation in the presence of species confers a benefit greater than any cost arising from increasing the complexity of the model, though we note covariate collinearity could still obfuscate any attempted model interpretation. Conversely the role of changing pseudo-absence patterns away from random did not repeat the results of previous work with thinning (Fourcade *et al*. 2014) and spatial weighting (Phillips *et al*. 2009) providing no clear gain in performance. The use of ‘pseudo-absence’ or ‘background’ points is simply a pragmatic solution to the common problem of the lack of known non-occupied sites (i.e. input data contains only the recorded presence or ‘sightings’ of target species).

Some recent developments have focussed instead on log-Gaussian Cox processes (Simpson *et al*. 2015) and Poisson point process (Renner & Warton 2013; Warton, Renner & Ramp 2013) which, instead of using presence/absence data, use the spatial density of presence points as the dependent variable in a regression. Alternatively, “range bagging” (Drake 2015) looks to bootstrap the variation in the multi-dimensional, environmental limits of a species, given random subsets of covariates and presence points. With these presence-only modelling approaches it is still unclear how they deal with non-biological, anthropogenic drivers of point density. For systematically collected data this may not be an issue, but for large citizen-science based or historic, museum-sourced databases unwanted spatial patterning of sample points may be a significant source of unresolved bias.

The R-INLA package appears to offer additional benefits beyond spatially-explicit modelling. The combination of using a complex spatial latent field to capture spatial processes and an underlying simpler ‘regression’ equation for the species’ relationship to environmental variables, means that (when compared to boosted regression trees and lasso techniques) the fixed effects are more straightforward to interpret (i.e. per unit change in x results in per unit change in y). Another benefit of a Bayesian approach is the capturing of uncertainty for each predicted value, with predictive uncertainty an often ignored aspect of SDM modelling and prediction. R-INLA models are extremely flexible in their specification, with spatial autocorrelation and observer bias being straightforwardly incorporated as random effects, while standard error “families” such as Gaussian, Poisson, binomial, and a variety of zero-inflated models, can be used interchangeably (Rue, Martino & Chopin 2009). This method, therefore, has a built-in potential for extending SDM analysis away from simple binomial model by, for example, incorporating two or more types of data (Warton *et al*. 2015) or hierarchical seasonal models (Redding *et al*. 2015). We hope that our study will aid the uptake of such fast spatial Bayesian methods, as this approach shows great promise for other analysis throughout ecology and evolutionary biology, especially in situations where non-independent samples are commonly experienced.

## Acknowledgements

We thank Anna Heath, Daniel Simpson and Sonia Tiedt for useful early discussions. This work, was funded with support from the Ecosystem Services for Poverty Alleviation Programme (ESPA), Dynamic Drivers of Disease in Africa Consortium, NERC project no. NE-J001570-1. The ESPA programme is funded by the Department for International Development (DFID), the Economic and Social Research Council (ESRC) and the Natural Environment Research Council (NERC).

## Data Accesibility

Simulated datasets and all code will be made available on Figshare.

## Supporting Information

Additional Supporting Information may be found in the online version of this article

